## Supplementary Information for "Convergent evolution of plant pattern recognition receptors sensing cysteine-rich patterns from three microbial kingdoms"

This file includes:
Extended data figures 1 to 10
Supplementary Tables S1 to S5

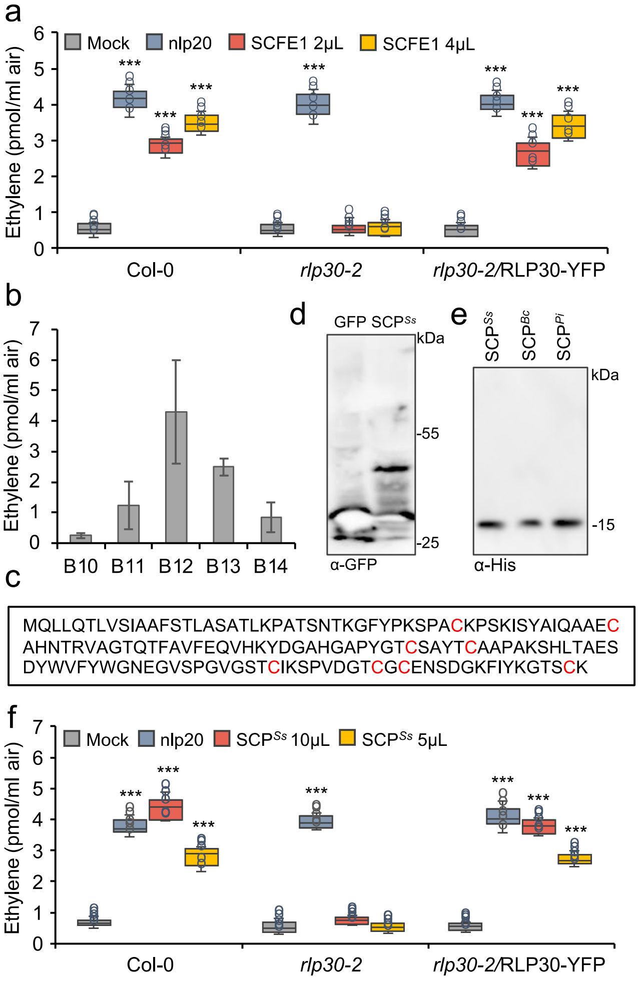


**Extended Fig. 1 Identification of an Arabidopsis defense-stimulating protein in the *S. sclerotiorum* SCFE1 preparation. a,** Ethylene accumulation in Col-0 wild-type plants, *rlp30-2* mutants or an *rlp30-2* line complemented with a *p35S::RL30-YFP* construct 4 h after treatment with water (mock), 1 µM nlp20, or 2 or 4 µl per 500 µl assay volume of a SCFE1 preparation. **b,** Ethylene accumulation in Col-0 wild-type plants treated with indicated SCFE1 fractions. **c,** Amino acid sequence of $SCP^{Ss}$ with the eight cysteine residues depicted in red. **d,e,** Western Blot analysis of SCP from *S. sclerotiorum* ($SCP^{Ss}$), *B. cinerea* ($SCP^{Bc}$), or *P. infestans* ($SCP^{Pi}$) produced in the *N. benthamiana* apoplast (**d**) or purified from *P. pastoris* (**e**) using anti-GFP or anti-His antibodies, respectively. **f,** Ethylene accumulation in Col-0 wild-type plants or *rlp30-2* mutants and complementation line 4 h after treatment with GFP purified from *N. benthamiana* apoplasts (mock), 1 µM nlp20, or given volumes per 500 µl assay volume of $SCP^{Ss}$ purified from *N. benthamiana* apoplasts. Data points are indicated as dots ($n$ = 6 for a; $n$ = 6 for e) and plotted as box plots (centre line, median; bounds of box, the first and third quartiles; whiskers, 1.5 times the interquartile range; error bar, minima and maxima). Statistically significant differences (**a,f**) from mock treatments in the respective plants are indicated (two-sided Student's t-test, ***$P \leq 0.001$). Each experiment was repeated three times with similar results.

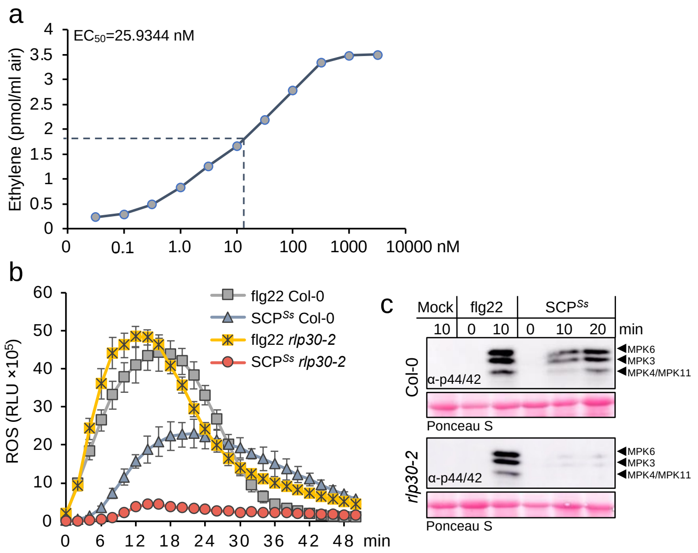


**Extended Fig. 2 Recombinant SCP$^{Ss}$ induces PTI responses. a,** Determination of $EC_{50}$ values using ethylene accumulation in *Arabidopsis* Col-0 wild-type plants after treatment with increasing concentrations of recombinant SCP$^{Ss}$ (produced in *Pichia*). $EC_{50}$ values and curve fit were calculated using $EC_{50}$ Calculator (https://www.aatbio.com/tools/ec50-calculator). **b,** ROS production in leaf pieces of *Arabidopsis* Col-0 wild-type plants or *rlp30-2* mutants treated with 0.1 µM flg22, or 2 µM SCP$^{Ss}$. Given are relative light units (RLU) ± SD (*n*=6). **c,** MAPK activation in *Arabidopsis* Col-0 wild-type plants or *rlp30-2* mutants treated for the times indicated with water (mock), 0.1 µM flg22, or 1 µM SCP$^{Ss}$. MAPK activation was detected by immunoblot using phospho-p44/p42 antibodies, equal loading was verified by staining of the membrane with Ponceau S Red. Experiments were repeated three times with similar results.

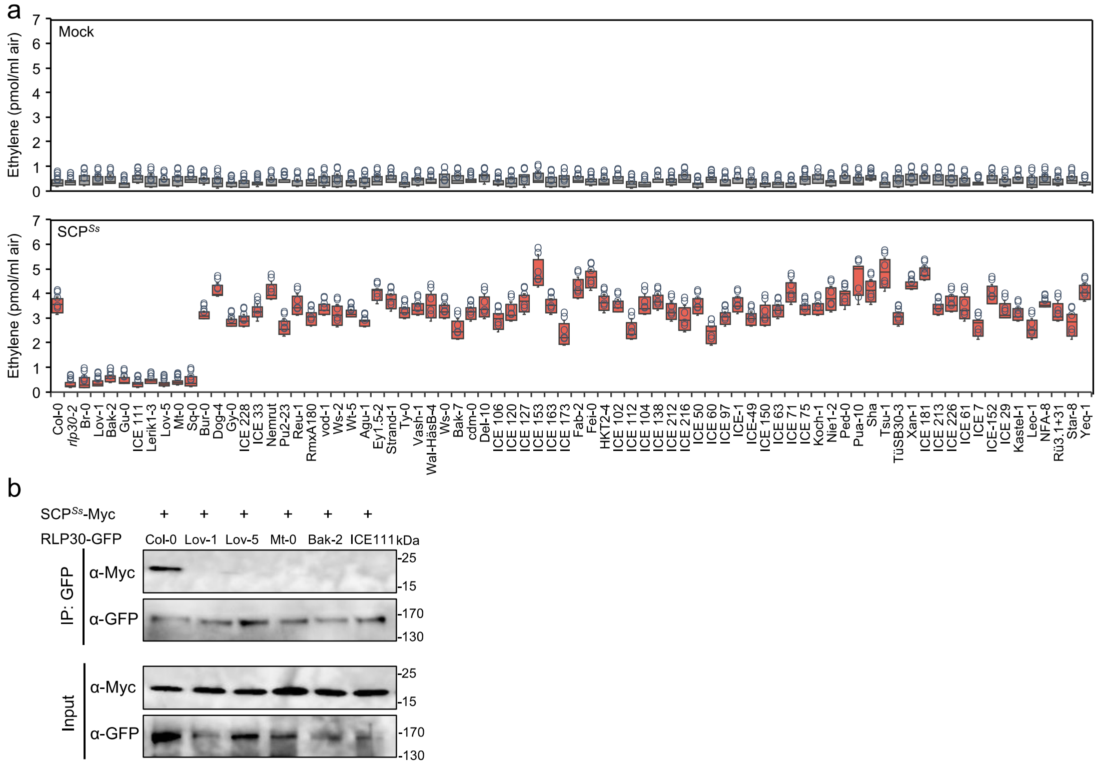


**Extended Fig. 3 SCP$^{Ss}$ binds to functional RLP30. a,** Natural variation in SCP$^{Ss}$ sensitivity among *Arabidopsis* accessions. 77 *Arabidopsis* accessions were tested for ethylene accumulation upon water (mock) or SCP$^{Ss}$ (1 µM) treatment. Data points are indicated as dots ($n$ = 3) and plotted as box plots (centre line, median; bounds of box, the first and third quartiles; whiskers, 1.5 times the interquartile range; error bar, minima and maxima). The experiment was repeated twice with similar results. **b,** Ligand-binding assay in *N. benthamiana* transiently expressing SCP$^{Ss}$-myc with RLP30-GFP from different *Arabidopsis* accessions. Proteins extracted from *N. benthamiana* leaves expressing indicated protein combinations (Input) were used for co-immunoprecipitation with GFP-trap beads (IP:GFP) and immunoblotting with tag-specific antibodies. The experiment was repeated three times with similar results.

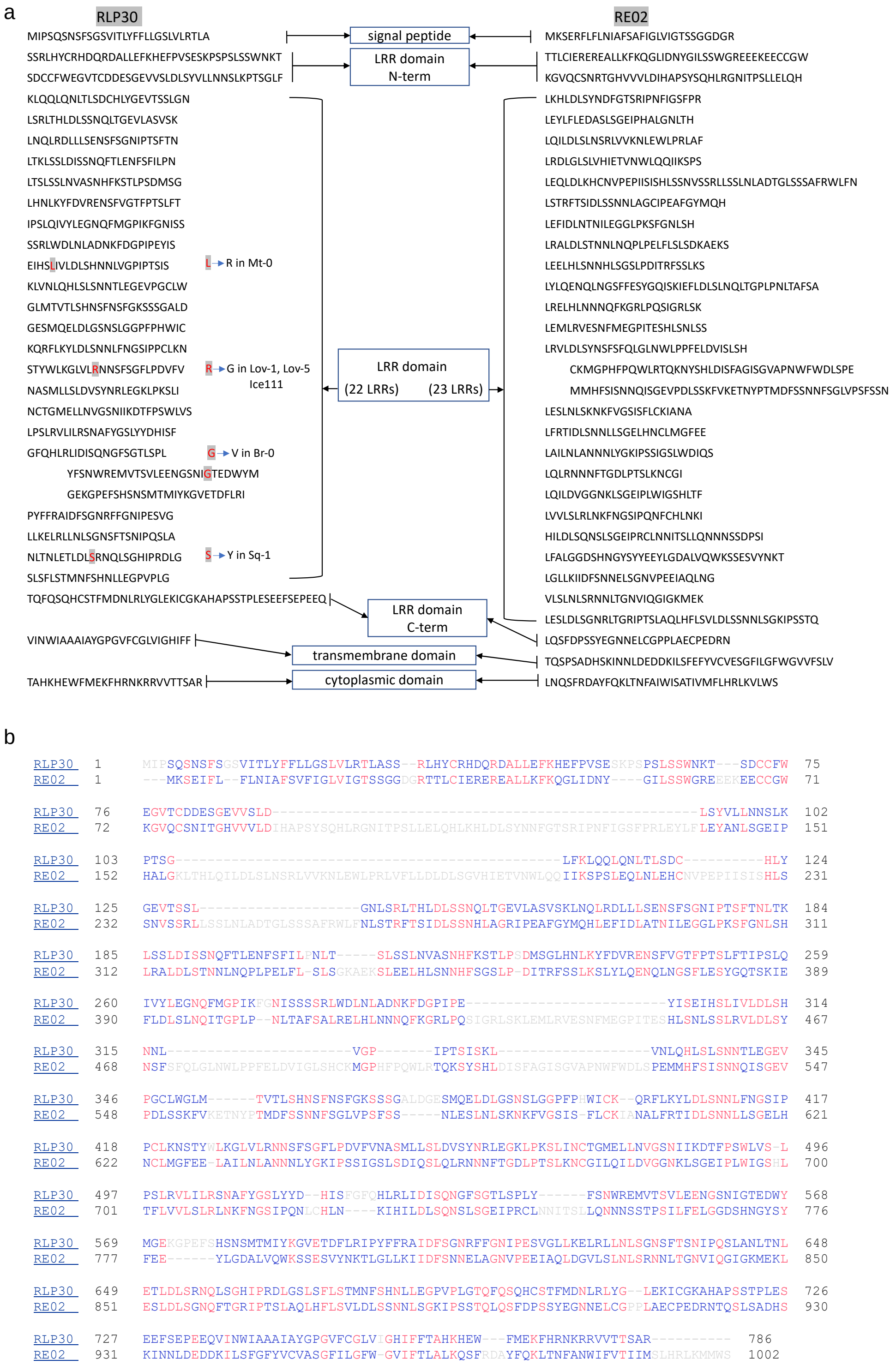


**Extended Fig. 4 Comparison of RLP30 and RE02 protein sequences. a,** Protein sequence display of RLP30 and RE02. Amino acids marked in red in the RLP30 sequence indicate amino acid changes in accessions that are insensitive to SCP$^{Ss}$. **b,** Protein sequence alignment performed by ClustalW, conserved residues between RLP30 and RE02 are shown in red.

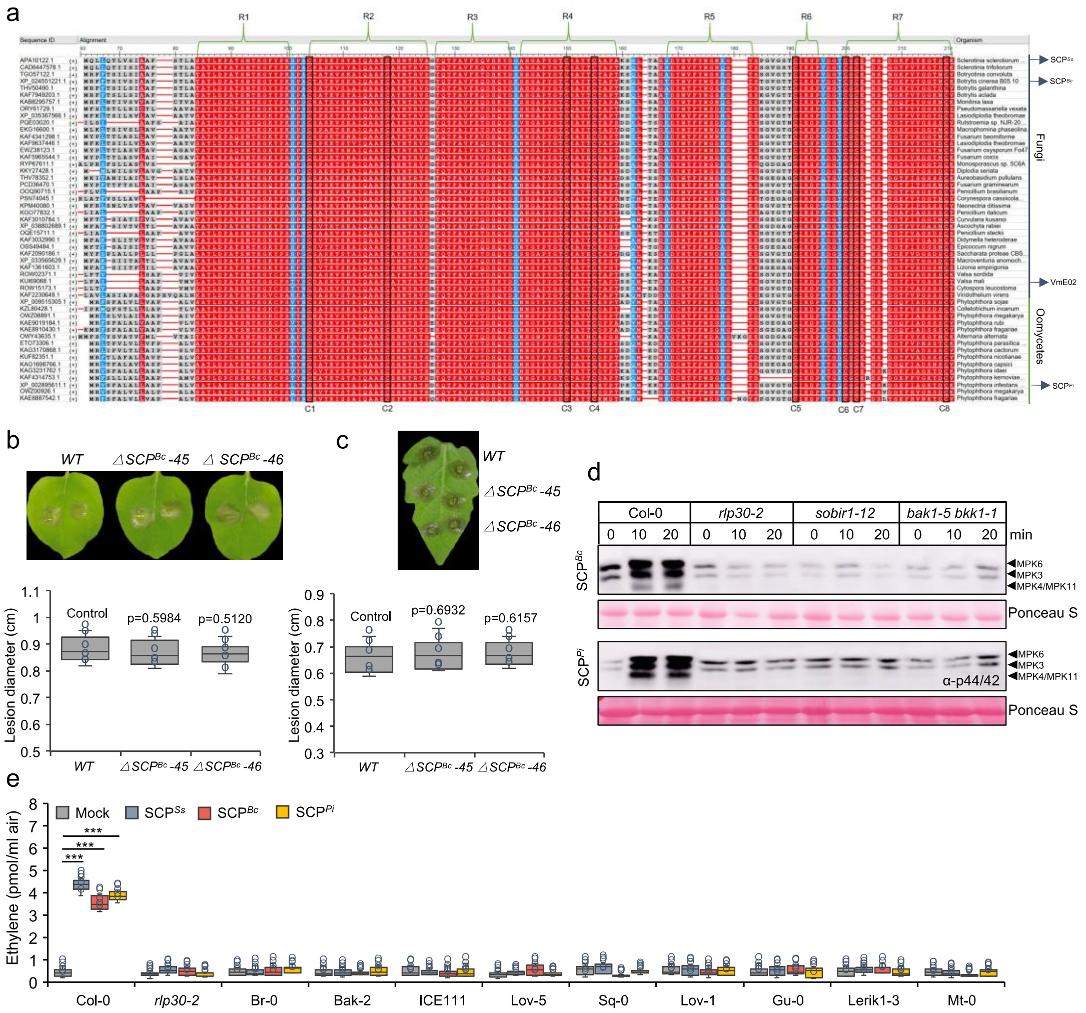


**Extended Fig. 5 $SCP^{Ss}$ is conserved in fungi and oomycetes. a,** Amino acid alignment of $SCP^{Ss}$ (top sequence) and its homologous sequences (Protein IDs given on the left) from the species indicated on the right. Conserved regions in red and conserved cysteine residues are numbered at the top (R1 to R7) and bottom (C1 to C7), respectively. **b,c,** Lesion formation of two independent *B. cinerea* depletion mutants of *$SCP^{Bc}$* (also named *Bcplp1*[1]) on *N. benthamiana* (**b**) or tomato (**c**) leaves 48 hours post inoculation and determination of lesion size. **d,** MAPK activation in *Arabidopsis* Col-0 wild-type plants or *rlp30-2, sobir1-12 and bak1-5 bkk1-1* mutants treated with 1 µM $SCP^{Bc}$ or $SCP^{Pi}$ for the times indicated. MAPK activation was detected by immunoblot using phospho-p44/p42 antibodies, equal loading of Ribulose-1,5-bisphosphate-carboxylase/oxygenase (Rubisco) was verified by staining of the membrane with Ponceau S Red. **e,** Ethylene accumulation in *Arabidopsis* wild-type plants (Col-0), or various $SCP^{Ss}$-insensitive accessions 4 h after treatment with water (mock), 1 µM $SCP^{Ss}$, $SCP^{Bc}$, or $SCP^{Pi}$. Data points are indicated as dots ($n$ = 6) and plotted as box plots (centre line, median; bounds of box, the first and third quartiles; whiskers, 1.5 times the interquartile range; error bar, minima and maxima). Statistically significant differences (**e**) from mock treatments in the respective plants are indicated (two-sided Student's t-test, ***$P \leq 0.001$). The experiments were repeated three times with similar results.

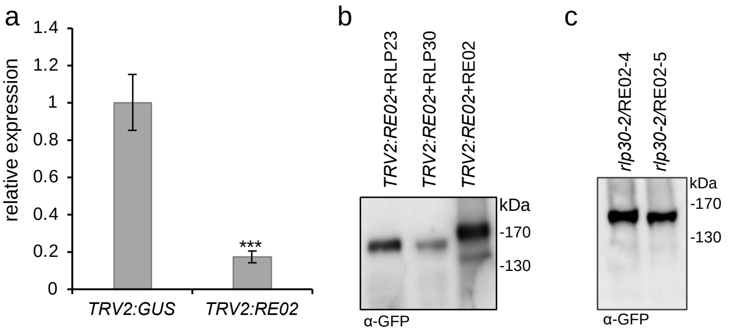


**Extended Fig. 6 Successful silencing of RE02 and heterologous complementation. a,** RT-qPCR analysis of relative expression of *RE02* in corresponding VIGS-silenced *N. benthamiana* leaves using gene-specific primers. Expression of *RE02* was normalized to the levels of *NbActin* transcript and is presented relative to the *TRV2:GUS* control which was set to 1. Statistically significant differences were determined using a two-sided Student's t-test (***$P \leq 0.001$). **b,** Western Blot analysis on protein extracts from *N. benthamiana* plants silenced for *RE02* and transiently expressing GFP-tagged RLP23, RLP30, or RE02 using an anti-GFP antibody. **c,** Western Blot analysis on protein extracts from two independent Arabidopsis *rlp30-2* lines stably expressing *p35S::RE02-GFP* using an anti-GFP antibody.

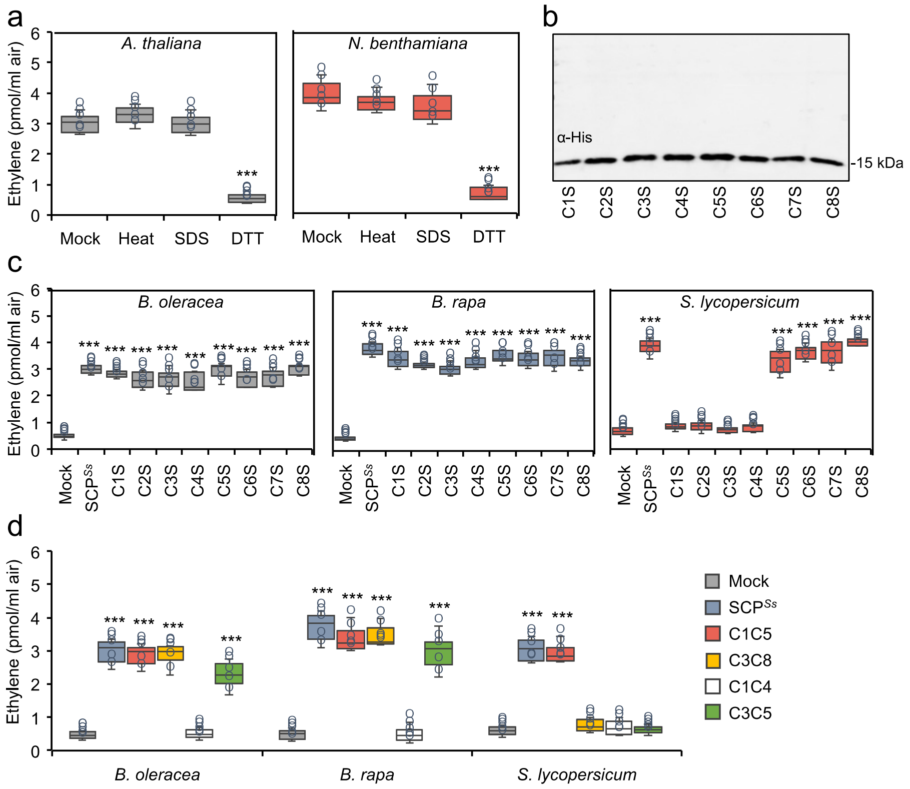


**Extended Fig. 7 Different requirements for cysteines in SCP$^{Ss}$ for its immunogenic activity in Brassicaceae and Solanaceae. a,** Ethylene accumulation in *A. thaliana* Col-0 or *N. benthamiana* plants treated with 1 µM water-treated SCP$^{Ss}$ (mock), or SCP$^{Ss}$ that was pre-treated for 1h at 95°C (heat), 1 % SDS, or 100 µM DTT. **b,** Western Blot analysis of SCP$^{Ss}$ with individual cysteine to serine mutations purified from *P. pastoris* using anti-His antiserum. **c,** Ethylene accumulation in *Brassica oleracea*, *B. rapa,* or *Solanum lycopersicum* plants after 4 h treatment with water (mock), SCP$^{Ss}$, and SCP$^{Ss}$ with individual cysteine to serine mutations. **d,** Ethylene accumulation in *B. oleracea*, *B. rapa,* or *S. lycopersicum* plants after 4 h treatment with water (mock), SCP$^{Ss}$, or SCP$^{Ss}$ truncations depicted in Fig. 3a. Data points (**a,c,d**) are indicated as dots ($n$ = 6) and plotted as box plots (centre line, median; bounds of box, the first and third quartiles; whiskers, 1.5 times the interquartile range; error bar, minima and maxima). Statistically significant differences from mock treatments in the respective plants are indicated (two-sided Student's t-test, ***$P \leq 0.001$). Each experiment was repeated three times with similar results.

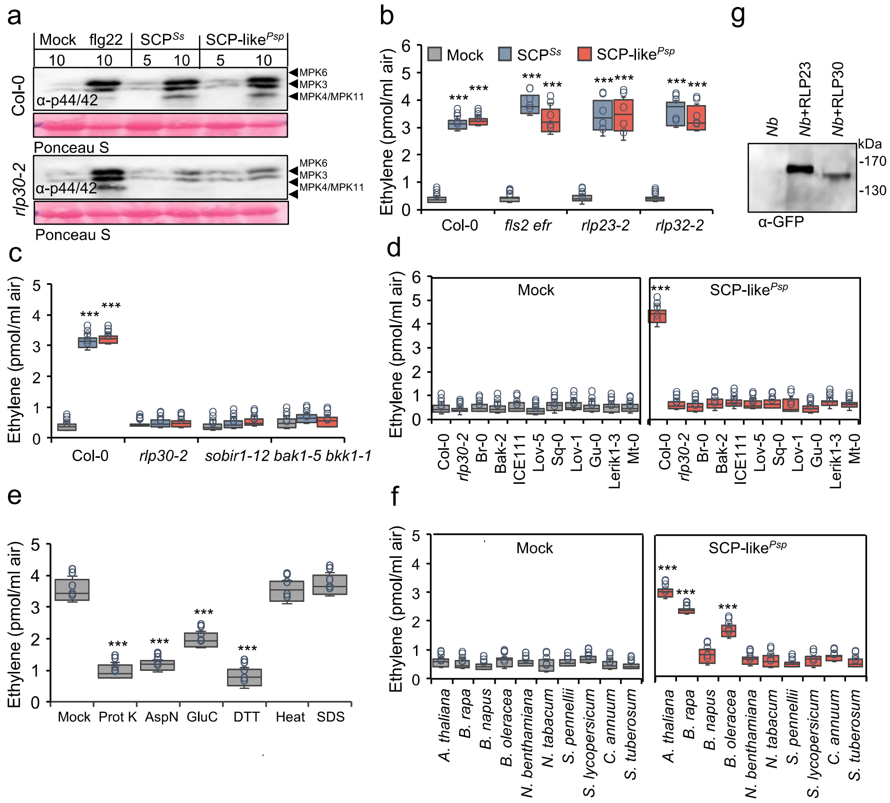


**Extended Fig. 8 *Pseudomonas* SCP-like induces RLP30-dependent PTI responses. a,** MAPK activation in Arabidopsis Col-0 wild-type plants or *rlp30-2* mutants infiltrated for the indicated times with 0.1 µM flg22, 1 µM SCP$^{Ss}$, or 1.5 µg/ml SCP-like$^{Psp}$. MAPK activation was detected by immunoblot using phospho-p44/p42 antibodies, equal loading was verified by staining of the membrane with Ponceau S Red. **b-d** Ethylene accumulation in Col-0 wild-type plants or indicated mutants (**b**,**c**) or SCP$^{Ss}$-insensitive accessions (**d**) 4 h after treatment with water (mock), 1 µM SCP$^{Ss}$, or 1.5 µg/ml SCP-like$^{Psp}$. **e,** Ethylene accumulation in Col-0 wild-type plants incubated for 4 h with water (mock), or 1.5 µg/ml SCP-like$^{Psp}$ treated for 4 h with 100 nM Proteinase K (Prot K), AspN, GluC, DTT, 1h at 95°C (heat), or 1 % SDS. **f,** Ethylene accumulation in Col-0 wild-type plants or indicated plants of the Brassicaceae and Solanaceae family 4 h after treatment with water (mock), or 1.5 µg/ml SCP-like$^{Psp}$. **g,** Western Blot analysis on protein extracts from *N. benthamiana* leaves transiently expressing RLP30-GFP or RLP23-GFP using an anti-GFP antibody. Data points (**b-f**) are indicated as dots ($n$ = 6) and plotted as box plots (centre line, median; bounds of box, the first and third quartiles; whiskers, 1.5 times the interquartile range; error bar, minima and maxima). Statistically significant differences from mock treatments in the respective plants are indicated (two-sided Student's t-test, $^{***}P \leq 0.001$). All experiments were repeated three times with similar results.

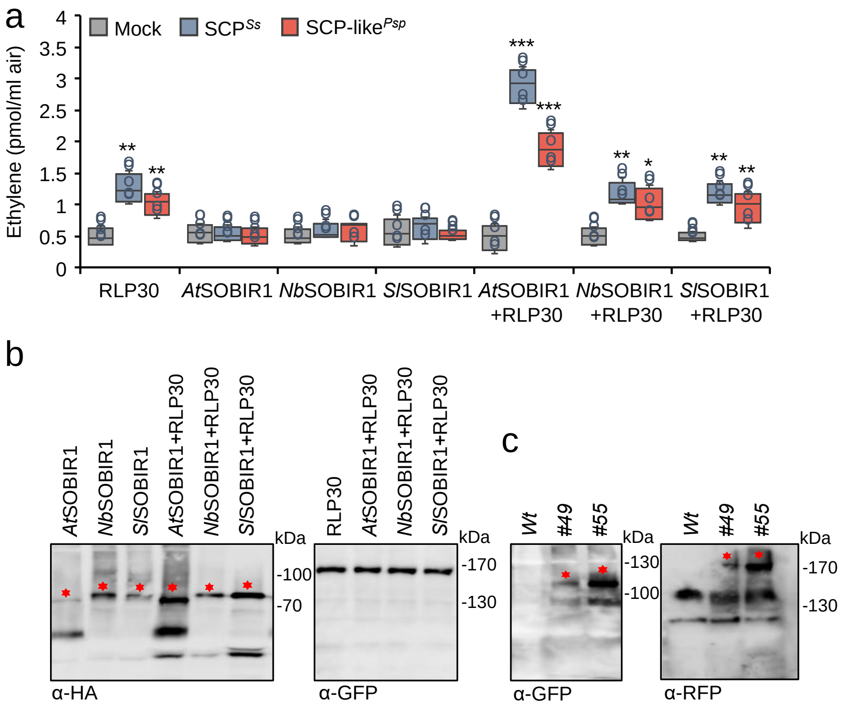


**Extended Fig. 9 AtSOBIR1 co-expression enhances RLP30 function in *N. tabacum*. a,** Ethylene accumulation in *N. tabacum* plants transiently expressing RLP30-GFP and/or SOBIR1 from Arabidopsis (*At*), *N. benthamiana* (*Nb*) *or S. lycopersicum* (*Sl*), either alone or in the indicated combination and treated for 4 h with water (mock), 1 µM SCP$^{Ss}$, or 1.5 µg/ml SCP-like$^{Psp}$. Data points are indicated as dots ($n$ = 6) and plotted as box plots (centre line, median; bounds of box, the first and third quartiles; whiskers, 1.5 times the interquartile range; error bar, minima and maxima). Statistically significant differences from mock treatments in the respective plants are indicated (two-sided Student's t-test, *$P \leq 0.05$, **$P \leq 0.01$, ***$P \leq 0.001$). **b,** Western Blot analysis with protein extracts from *N. tabacum* leaves transiently expressing RLP30-GFP and/or AtSOBIR1-HA, NbSOBIR1-HA or SlSOBIR1-HA as shown in (**a**) using an anti-GFP antibody for RLP30 detection and an anti-HA antibody for SOBIR1 detection. **c,** Western Blot analysis with protein extracts from two independent *N. tabacum* lines (*#49* and *#55*) stably expressing RLP30-RFP and AtSOBIR1-GFP using tag-specific antisera. The asterisks indicate the position of epitope-tagged proteins. All experiments were repeated three times with similar results.

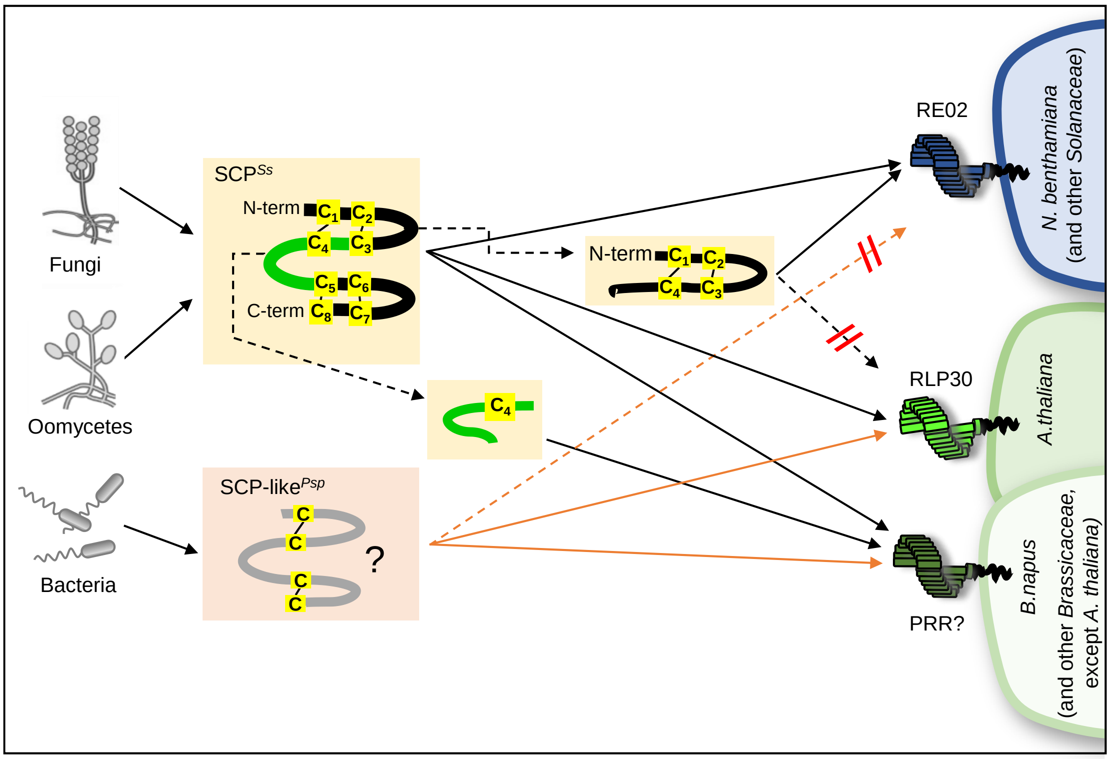


**Extended Fig. 10 RLP30 and RE02 mediate pattern recognition.** RLP30 recognizes SCP$^{Ss}$ and its homologs from different fungi and oomycetes, as well as an SCP$^{Ss}$-unrelated and conserved pattern from *Pseudomonas* (SCP-like$^{Psp}$). RE02, an LRR-RLP non-homologous to RLP30, mediates SCP$^{Ss}$ recognition in *N. benthamiana*. Unlike RLP30-mediated recognition of entire SCP$^{Ss}$ in *A. thaliana*, Brassica species and Solanaceae perceive a small immunogenic epitope and a disulfide-bond containing peptide of SCP$^{Ss}$, respectively.

### Supplementary Table S1. Candidate proteins within SCFE1 obtained by MS/MS-analyses

| Protein ID | detected unique peptides per protein | Sequence coverage [%] | Mol. weight [kDa] | Sequence length [amino acid count] | signal peptide present | cysteines | Cysteine content [cysteines/seq.length in %] | MS intensity (iBAQ) in SCFE1 fraction B9 | MS intensity (iBAQ) in SCFE1 fraction B11 | MS intensity (iBAQ) in SCFE1 fraction B12 | MS intensity (iBAQ) in SCFE1 fraction B13 | MS intensity (iBAQ) in SCFE1 fraction B15 |
|---|---|---|---|---|---|---|---|---|---|---|---|---|
| Sscle06g048920 | 2 | 17 | 15,481 | 147 | yes | 8 | 5,4 | 0 | 3349300 | 10808000 | 10764000 | 0 |
| Sscle10g077860 | 2 | 18,8 | 21,712 | 191 | no | 1 | 0,5 | 0 | 1958400 | 626520 | 1260200 | 0 |
| Sscle12g090490 | 2 | 6,5 | 26,041 | 245 | yes | 5 | 2 | 47343000 | 273220000 | 102210000 | 116290000 | 0 |
| Sscle06g050580 | 2 | 13,4 | 26,089 | 238 | yes | 4 | 1,7 | 0 | 0 | 270780 | 1350500 | 0 |
| Sscle01g009200 | 2 | 12,1 | 28,531 | 273 | yes | 5 | 1,8 | 769810 | 178500 | 0 | 0 | 638100 |
| Sscle16g108160 | 6 | 19,9 | 34,201 | 327 | yes | 3 | 0,9 | 790000000 | 584600000 | 326300000 | 376090000 | 13573000 |
| Sscle09g073350 | 2 | 5,3 | 37,229 | 339 | yes | 10 | 2,9 | 0 | 1079300 | 113530 | 267700 | 0 |
| Sscle16g108170 | 3 | 3,7 | 37,672 | 380 | yes | 8 | 2,1 | 0 | 121680000 | 5940700 | 4476100 | 810830 |
| Sscle10g080050 | 6 | 12,6 | 40,184 | 358 | yes | 4 | 1,1 | 525860 | 162740000 | 384270000 | 1745000000 | 730920000 |
| Sscle14g099090 | 5 | 16,5 | 41,639 | 375 | no | 4 | 1,1 | 977170 | 1847300 | 1657100 | 1846100 | 3282500 |
| Sscle03g030530 | 8 | 21,9 | 42,399 | 397 | no | 3 | 0,8 | 290420 | 303070000 | 151640000 | 695190000 | 24978000 |
| Sscle04g040020 | 2 | 4,8 | 48,544 | 475 | yes | 11 | 2,3 | 0 | 3365600 | 2651900 | 4968400 | 0 |
| Sscle03g028450 | 9 | 17,3 | 49,123 | 445 | yes | 4 | 0,9 | 0 | 4928200 | 6765900 | 19093000 | 0 |
| Sscle02g017490 | 2 | 1,8 | 62,841 | 605 | yes | 8 | 1,3 | 0 | 747520 | 879350 | 2244900 | 0 |
| Sscle11g085960 | 2 | 5,2 | 66,699 | 611 | yes | 6 | 1 | 0 | 6739000 | 0 | 10596000 | 0 |
| Sscle10g080270 | 19 | 30,2 | 67,026 | 616 | yes | 10 | 1,6 | 12583000 | 245180000 | 423460000 | 570200000 | 16638000 |
| Sscle08g063080 | 13 | 23,9 | 67,41 | 610 | yes | 8 | 1,3 | 28634000 | 109240000 | 61576000 | 80709000 | 6074600 |
| Sscle15g106280 | 3 | 6,2 | 68,529 | 632 | yes | 6 | 0,9 | 583570 | 1179100 | 1547600 | 4099900 | 490000 |
| Sscle14g100060 | 2 | 4,9 | 69,742 | 613 | no | 9 | 1,5 | 104900 | 556270 | 943020 | 2439600 | 1364000 |
| Sscle06g050510 | 3 | 6,7 | 72,965 | 669 | yes | 9 | 1,3 | 0 | 242100 | 0 | 1844300 | 1419800 |
| Sscle07g057170 | 2 | 5,2 | 99,173 | 907 | yes | 12 | 1,3 | 0 | 36568 | 0 | 370790 | 0 |
| Sscle09g074570 | 4 | 4,2 | 109,64 | 1012 | yes | 2 | 0,2 | 0 | 1325600 | 1280500 | 1194200 | 0 |

**Supplementary Table S2. *Arabidopsis* lines used in this study**

| Line | Locus | Description | Reference |
|---|---|---|---|
| *bak1-5 bkk1-1* | At4g33430<br>At2g13790 | Double mutant of bak1-5 and SALK_044334 | [2] |
| *fls2 efr* | At5g46330<br>At5g20480 | Double mutant of SAIL_691_C4 and SALK_044334 | [3] |
| *rlp23-1* | At2g32680 | Insertion, SALK_034225 | [4] |
| *rlp30-2* | At3g05360 | Insertion, SALK_008911 | [5] |
| *rlp30-2/RLP30-YFP* | At3g05360 | Insertion, SALK_008911, complemented with YFP-tagged RLP30 | This study |
| *rlp30-2/NbRE02-GFP* | At3g05360 | Insertion, SALK_008911, complemented with GFP-tagged NbRE02 | This study |
| *rlp32-2* | At3g05650 | Insertion, SM_3_33092 | [6] |
| *sobir1-12* | At2g31880 | Insertion, SALK_050715, *sobir1-12* | [7] |

**Supplementary Table S3. Primers used for cloning**

| Template | Expression in | Primer name | Primer sequence (5' – 3') |
|---|---|---|---|
| *NbRE02* | *rlp30-2*<br>*N. tabacum,*<br>*N. benthamiana* | B_RE02-F | tatggtctcatctgaacaATGAAAAGTGAGAGATTT |
| | | D_RE02-R | ttggtctctccttACTCCAGAGCACCTTCAATCTGTG |
| *NbRE02* | *VIGS,*<br>*N. benthamiana* | TRV2:NbRE02_F | GTGAGCTCGGTACCGGATCCGAACTCCCGTCTA GTAGT |
| | | TRV2:NbRE02_R | TGAGTAAGGTTACCGAATTCACCTTCTAGAATATT TGTATT |
| *RLP30, all accessions* | *N. tabacum,*<br>*N. benthamiana* | RLP30-F | ATGATTCCAAGCCAATCTAATTCC |
| | | RLP30-R(noStop) | ACGAGCACTTGTGGTGACTAC |
| *SCP$^{Ss}$* | *N. benthamiana* apoplast | SCP-N_F | CATTTACGAACGATAGGGTACCCCCATGCAACTC CTCCAAACCC |
| | | SCP-N_R | TGCTCACCATGGATCCGTCGACCCCTTTACAAG AAGTCCCCTTGTAGATAAAC |
| *SCP$^{Ss}$*<br>*SCP$^{Ss}$*(C1-7-S) | *Pichia pastoris* | SCP-P_F | CGGAATTCATGACCCTCAAACCCGCTACCTC |
| | | SCP-P_R | GCTCTAGACCTTTACAAGAAGTCCCCTTGTAGAT AAACTTACC |
| *SCP$^{Ss}$*(C8-S) | *Pichia pastoris* | SCP$^{C8-S}$-P_F | GCGGAATTCATGACCCTCAAACCCGCTACCTC |
| | | SCP$^{C8-S}$-P_R | GCTCTAGACCTTTAGAAGAAGTCCCCTTGTAGAT AAACTTACC |
| *SCP$^{Bc}$* | *Pichia pastoris* | SCP$^{Bc}$-P_F | CGGAATTCATGCTCGACCCCGCTACCTCAAAC |
| | | SCP$^{Bc}$-P_R | GCTCTAGACCCTTGCAGTTCGTTCCTCCATAAAC AAAC |
| *SCP$^{Pi}$* | *Pichia pastoris* | SCP$^{Pi}$-P_F | CGGAATTCATGGCTCCTTGCCGCACCAATAG |
| | | SCP$^{Pi}$-P_R | GCTCTAGACCTTTGCAGTCCGTCTTGCCG |
| *SCP$^{Ss}$*(C1C5) | *Pichia pastoris* | SCP$^{C1C5}$P_F | CGGAATTCATGACCCTCAAACCCGCTACCTC |
| | | SCP$^{C1C5}$P_R | GCTCTAGACCAATACAAGTCGAGCCCACACCA |
| *SCP$^{Ss}$*(C3C8) | *Pichia pastoris* | SCP$^{C3C8}$P_F | CGGAATTCATGACTTGTTCCGCTTATACCTGTGC T |
| | | SCP$^{C3C8}$P_R | GCTCTAGACCTTTACAAGAAGTCCCCTTGTAGAT AAACTTACC |
| *SCP$^{Ss}$*(C1C4) | *Pichia pastoris* | SCP$^{C1C4}$P_F | CGGAATTCATGGCTTGCAAACCAAGCAAAATCT CC |
| | | SCP$^{C1C4}$P_R | GCTCTAGACCAGCACAGGTATAAGCGGAACAAG |
| *SOBIR1* | *N. tabacum,*<br>*N. benthamiana* | SOBIR1-F | ATGGCTGTTCCCACGGGAAG |
| | | SOBIR1-R(noStop) | GTGCTTGATCTGGGACAACATG |

**Supplementary Table S4. Primers used for quantitative RT-PCR (VIGS)**

| Gene | Primer name | Primer sequence (5' – 3') |
|---|---|---|
| *NbActin* | NbActin_F | TGGTCGTACCACCGGTATTGTGTT |
| | NbActin_R | TCACTTGCCCATCAGGAAGCTCAT |
| *NbRE02* | qRT-NbRE02_F | TGCATCCCTGAAGCCTTTGG |
| | qRT-NbRE02_R | TCAGGAAGTGGTTGGTTCAAGT |

**Supplementary Table S5. SCP1 gBlocks used as PCR-templates for *Pichia* expression**

Codons for cysteines are in red, point mutations are highlighted in blue

| Name | Synthetic gBlock (cysteine codons in red, mutations in blue and highlighted) |
|---|---|
| C1S | ACCCTCAAACCCGCTACCTCAAACACAAAAGGCTTCTACCCCAAATCTCCAGCTTCCAAACCAAGC<br>AAAATCTCCTACGCCATCCAAGCCGCCGAATGCGCCCACAACACCCGTGTAGCCGGCACGCAAAC<br>CTTCGCCGTCTTCGAACAAGTCCACAAATATGATGGCGCACACGGTGCTCCCTATGGAACTTGTTC<br>CGCTTATACCTGTGCTGCGCCTGCGAAATCACACTTGACGGCTGAGTCTGATTATTGGGTGTTTTAT<br>TGGGGTAATGAGGGGGTTAGTCCTGGTGTGGGCTCGACTTGTATTAAGAGTCCTGTGGATGGGAC<br>TTGTGGGTGTGAGAATTCGGATGGTAAGTTTATCTACAAGGGGACTTCTTGTAAA |
| C2S | ACCCTCAAACCCGCTACCTCAAACACAAAAGGCTTCTACCCCAAATCTCCAGCTTGCAAACCAAGC<br>AAAATCTCCTACGCCATCCAAGCCGCCGAATCCGCCCACAACACCCGTGTAGCCGGCACGCAAAC<br>CTTCGCCGTCTTCGAACAAGTCCACAAATATGATGGCGCACACGGTGCTCCCTATGGAACTTGTTC<br>CGCTTATACCTGTGCTGCGCCTGCGAAATCACACTTGACGGCTGAGTCTGATTATTGGGTGTTTTAT<br>TGGGGTAATGAGGGGGTTAGTCCTGGTGTGGGCTCGACTTGTATTAAGAGTCCTGTGGATGGGAC<br>TTGTGGGTGTGAGAATTCGGATGGTAAGTTTATCTACAAGGGGACTTCTTGTAAA |
| C3S | ACCCTCAAACCCGCTACCTCAAACACAAAAGGCTTCTACCCCAAATCTCCAGCTTGCAAACCAAGC<br>AAAATCTCCTACGCCATCCAAGCCGCCGAATGCGCCCACAACACCCGTGTAGCCGGCACGCAAAC<br>CTTCGCCGTCTTCGAACAAGTCCACAAATATGATGGCGCACACGGTGCTCCCTATGGAACTTCTTC<br>CGCTTATACCTGTGCTGCGCCTGCGAAATCACACTTGACGGCTGAGTCTGATTATTGGGTGTTTTAT<br>TGGGGTAATGAGGGGGTTAGTCCTGGTGTGGGCTCGACTTGTATTAAGAGTCCTGTGGATGGGAC<br>TTGTGGGTGTGAGAATTCGGATGGTAAGTTTATCTACAAGGGGACTTCTTGTAAA |
| C4S | ACCCTCAAACCCGCTACCTCAAACACAAAAGGCTTCTACCCCAAATCTCCAGCTTGCAAACCAAGC<br>AAAATCTCCTACGCCATCCAAGCCGCCGAATGCGCCCACAACACCCGTGTAGCCGGCACGCAAAC<br>CTTCGCCGTCTTCGAACAAGTCCACAAATATGATGGCGCACACGGTGCTCCCTATGGAACTTGTTC<br>CGCTTATACCTCTGCTGCGCCTGCGAAATCACACTTGACGGCTGAGTCTGATTATTGGGTGTTTTAT<br>TGGGGTAATGAGGGGGTTAGTCCTGGTGTGGGCTCGACTTGTATTAAGAGTCCTGTGGATGGGAC<br>TTGTGGGTGTGAGAATTCGGATGGTAAGTTTATCTACAAGGGGACTTCTTGTAAA |
| C5S | ACCCTCAAACCCGCTACCTCAAACACAAAAGGCTTCTACCCCAAATCTCCAGCTTGCAAACCAAGC<br>AAAATCTCCTACGCCATCCAAGCCGCCGAATGCGCCCACAACACCCGTGTAGCCGGCACGCAAAC<br>CTTCGCCGTCTTCGAACAAGTCCACAAATATGATGGCGCACACGGTGCTCCCTATGGAACTTGTTC<br>CGCTTATACCTGTGCTGCGCCTGCGAAATCACACTTGACGGCTGAGTCTGATTATTGGGTGTTTTAT<br>TGGGGTAATGAGGGGGTTAGTCCTGGTGTGGGCTCGACTTCTATTAAGAGTCCTGTGGATGGGAC<br>TTGTGGGTGTGAGAATTCGGATGGTAAGTTTATCTACAAGGGGACTTCTTGTAAA |
| C6S | ACCCTCAAACCCGCTACCTCAAACACAAAAGGCTTCTACCCCAAATCTCCAGCTTGCAAACCAAGC<br>AAAATCTCCTACGCCATCCAAGCCGCCGAATGCGCCCACAACACCCGTGTAGCCGGCACGCAAAC<br>CTTCGCCGTCTTCGAACAAGTCCACAAATATGATGGCGCACACGGTGCTCCCTATGGAACTTGTTC<br>CGCTTATACCTGTGCTGCGCCTGCGAAATCACACTTGACGGCTGAGTCTGATTATTGGGTGTTTTAT<br>TGGGGTAATGAGGGGGTTAGTCCTGGTGTGGGCTCGACTTGTATTAAGAGTCCTGTGGATGGGAC<br>TTCTGGGTGTGAGAATTCGGATGGTAAGTTTATCTACAAGGGGACTTCTTGTAAA |
| C7S | ACCCTCAAACCCGCTACCTCAAACACAAAAGGCTTCTACCCCAAATCTCCAGCTTGCAAACCAAGC<br>AAAATCTCCTACGCCATCCAAGCCGCCGAATGCGCCCACAACACCCGTGTAGCCGGCACGCAAAC<br>CTTCGCCGTCTTCGAACAAGTCCACAAATATGATGGCGCACACGGTGCTCCCTATGGAACTTGTTC<br>CGCTTATACCTGTGCTGCGCCTGCGAAATCACACTTGACGGCTGAGTCTGATTATTGGGTGTTTTAT<br>TGGGGTAATGAGGGGGTTAGTCCTGGTGTGGGCTCGACTTGTATTAAGAGTCCTGTGGATGGGAC<br>TTGTGGGTCTGAGAATTCGGATGGTAAGTTTATCTACAAGGGGACTTCTTGTAAA |
| C8S | ACCCTCAAACCCGCTACCTCAAACACAAAAGGCTTCTACCCCAAATCTCCAGCTTGCAAACCAAGC<br>AAAATCTCCTACGCCATCCAAGCCGCCGAATGCGCCCACAACACCCGTGTAGCCGGCACGCAAAC<br>CTTCGCCGTCTTCGAACAAGTCCACAAATATGATGGCGCACACGGTGCTCCCTATGGAACTTGTTC<br>CGCTTATACCTGTGCTGCGCCTGCGAAATCACACTTGACGGCTGAGTCTGATTATTGGGTGTTTTAT<br>TGGGGTAATGAGGGGGTTAGTCCTGGTGTGGGCTCGACTTGTATTAAGAGTCCTGTGGATGGGAC<br>TTGTGGGTGTGAGAATTCGGATGGTAAGTTTATCTACAAGGGGACTTCTTCTAAA |
